# 3D scanning measurement method for seed size research

**DOI:** 10.1101/2024.08.22.609234

**Authors:** Hajnalka Málik-Roffa, Tamás Málik, Dávid Tőzsér, Béla Tóthmérész

**Author notes:** Correspondence: Hajnalka Málik-Roffa.

## Abstract

**Premise:** Traditionally used seed size and surface area calculation techniques are often inaccurate or destructive. This is especially relevant for organisms with irregular shapes. It is believed that three-dimensional (3D) printing and scanning could also be disruptive technologies. However, these methods hold immense potential for ecology and evolution sciences, which have a longstanding tradition of inventing and creating objects for research, education, and outreach.

**Methods and Results:** We describe how to 3D scan seeds reproducible on an inexpensive structured light based commodity 3D scanner. We then compared novel 3D and traditional 2D approaches by reproducibility and timing.

**Conclusions:** This study demonstrates how 3D scanning methods are easily applicable to the quantification of seed shape and providing an easy entry into it.

## 1. INTRODUCTION

Anticipating how ecosystems will react to rapid environmental changes has emerged as a great challenge in modern biology. Ecologists and evolutionary biologists tend to solve these challenges with Do-it-yourself and crafting skills. These solutions are manifold and usually customized to a specific application or problem, causing a considerable variation in model systems in ecology. A relative newcomer is likely to become a valuable tool in the ecologist toolbox: three-dimensional (3D) printing (Behm et al., 2018; Schtickzelle et al., 2020) and scanning (Reichert et al., 2016; Quigley, 2020).

To merge the physical and digital, 3D scanning is the way to create a digital 3D model of a real object by analyzing it or its surrounding environment to capture three-dimensional data about its shape and, preferably, its visual attributes (such as color). This technology enables precise digital representation of real-world objects, which creates many possibilities in fields like manufacturing or architecture.

3D scanning employs two main methods to generate a digital 3D model: either by using a scanner or a camera (photogrammetry). 3D scanners capture individual point locations of an object’s entire volume, which are then combined to form a 3D mesh. The location of each individual point defined in our case by a high-precision Blue Light which is a version of structured light scanning technologies (Cui et al., 2021; Kamal et al., 2014). It can capture a higher density of point clouds than laser scanning. For data collection a projector, camera, and lens system are combined. The projector creates typically straight lines light patterns on the object, which is then captured by the camera. The camera and projector are positioned at an angle to each other so the distance of every point can be calculated in the field of view. To capture an object’s entire volume, the scanner must either rotate around the object or the object must be rotated on a turntable in front of the scanner.

The photogrammetry approach creates a 3D mesh based on the analysis of a series of photographic images shot from various directions (Mesken et al., 2022; Ferrari et al., 2021). When it comes to scanning small objects, using a scanner is a reliable choice (Cui et al., 2021; Ma et al., 2009).

The major goal of this article is to describe a general workflow for 3D scanning seeds or any irregularly shaped ecological objects with high efficiency, show that it has a similar reproducibility rate to traditional measuring methods, and show the timing difference between the two methods. Also, promote usage in other areas of ecology where these tools are not utilized yet.

## 2. METHODS AND RESULTS

### 2.1 Species

During this project, fifteen pieces of Quercus robur (Fagaceae) and Aesculus hippocastanum (Sapindaceae) seeds were scanned. All of them were marked with a number for easier identification.

### 2.2 3D scanner selection

For this study, we chose the Revopoint Mini (Revopoint 2022) 3D scanner which are controlled by the Revo Scan 5 software. With scan speed up to 10fps (frames per second), short scan time can be achieved with precision up to 0.02mm and accuracy up to 0.05mm. It is an inexpensive commodity hardware for general use with sufficient precision and accuracy. Many previous studies used industrial scanners which naturally have better performance (Reichert et al., 2016; Laura et al., 2009). Therefore, this study aimed to reveal whether the use of such industrial tools can be replaced – in most of the cases – by using one of the best value for money 3D scanners on the market.

### 2.3 3D scanning and modeling

*Aesculus hippocastanum* seeds are nut-brown with glossy surface, the conditions where scanning with structured light technics is the most inaccurate. The following step is probably skippable for objects with matt finish or brighter colors. To achieve efficient object boundary recognition, two or three layers of scan helper spray (later scanning spray) were applied on each seed, followed by 3-5 minutes of drying. Two scanning sprays were tested in advance: ATTBLIME ABP and ATTBLIME AB ZERO. The difference between these two was found in persistence: the AB ZERO stays on the surface for up to one hour, followed by a sublimation and dissolution in 1-2 hours, while the ABP stays permanently until it is cleaned. Waiting at least one day after use is recommended for easier cleaning. We noticed no difference in the effectiveness of the sprays, but mostly, ABP was used. *Quercus robur* seeds are less challenging to scan, but for optimal results, we also covered them with ABP spray. Seeds were pierced with a needle in the middle and placed one by one on a small hard sponge, which was fixed with double side tape to the dual-axis rotating plate. For light treatment, we used a Puluz Photo box (24 × 23 × 23 cm) to ensure a solid background and constant light exposure (Figure 1).

**Figure 1.**
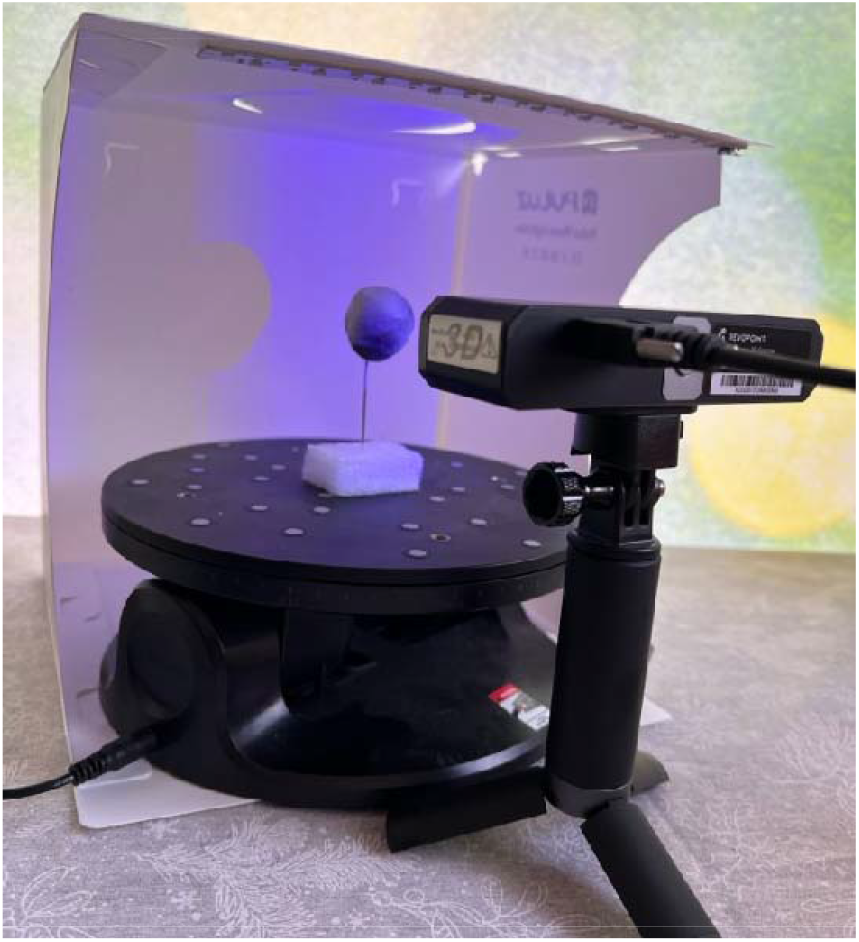
The scanning setup

After preparation, the Scans (accuracy: High Accuracy, tracking mode: Feature Tracking, Object Type: General Object, color scanning: off) were captured within 120–150s, in two to three complete rotations from one lower angle and two other from an another higher angle, commonly from -30° and 30° in respect to the scanned fragment. Scans, where the object tracking has lost, were repeated to avoid unnecessary work of ghost point removal.

### 2.4 3D model processing

Models were processed using an Apple Macbook Pro M1 14-inch 2021 laptop. After the point clouds were gathered, the points were fused with the ‘Advanced’ setting and with 0.15-0.18mm point distance based on the default settings of the Revo Scan 5 software for each model. The software has capabilities for post-processing the fused then 3D meshed models, so we intended to do as much work in a single software as possible. After fusing, a 3D mesh of the object has been generated with a grid size 0.15-0.20mm setting based on the software recommendation. The automatic hole filling was turned off during the meshing process. Even with the most careful handling, the layers of scanning sprays could be damaged, and small holes could be visible on the meshed surface. If a hole was less than 0.2mm, we used the built-in ‘Fill Holes’ function (methods: Curved) to correct it; otherwise, the scan was repeated. In this function, we used the automatic detection option rather than manually selecting which ones to fill.

As a final step before exporting the result model, all non-seed structures were edited out manually, such as the stand or the needle using the polygon or rectangular selection then the eraser (Figure 2).

**Figure 2.**
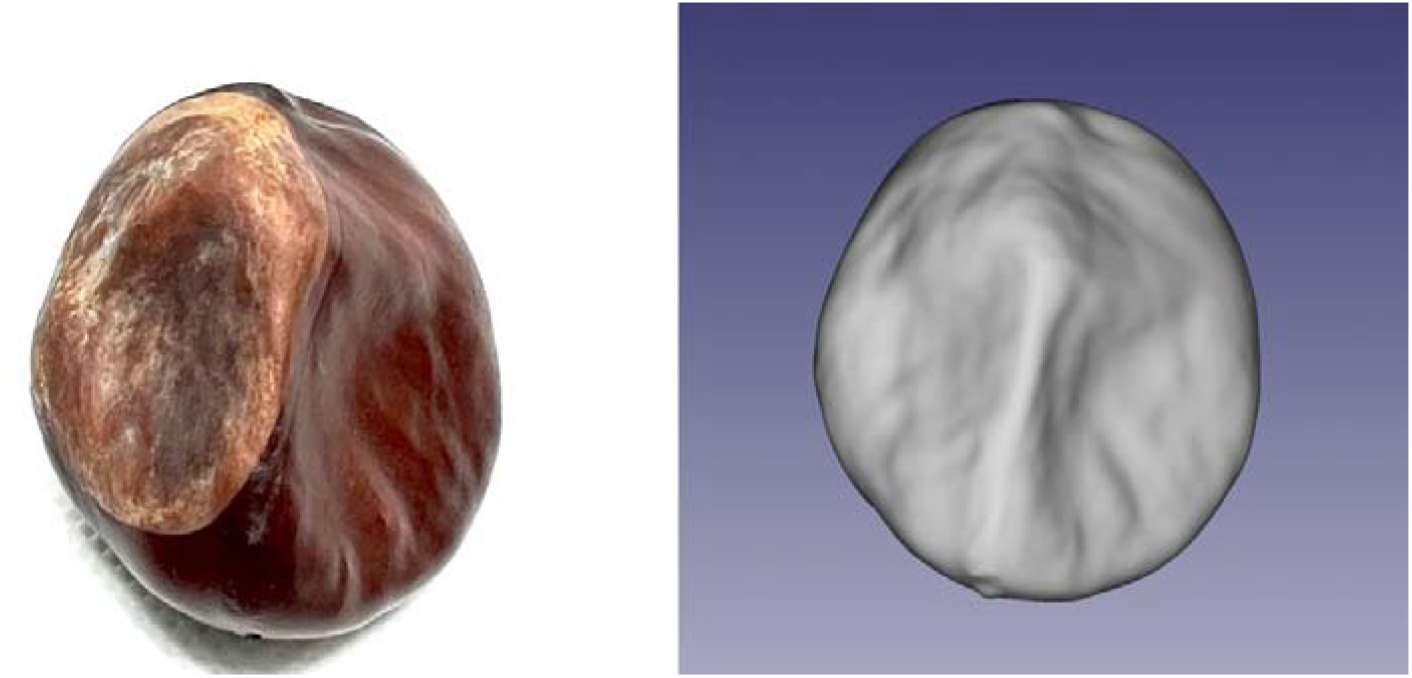
An Aesculus hippocastanum seed and its 3D-scanned model

### 2.5 3D computational measurement

For the analyses, meshes were exported to Wavefront “.obj” format and imported into FreeCAD (v0.21.1). Length and width were calculated via the FCInfo (v1.28c) macro addon (Figure 3).

**Figure 3.**
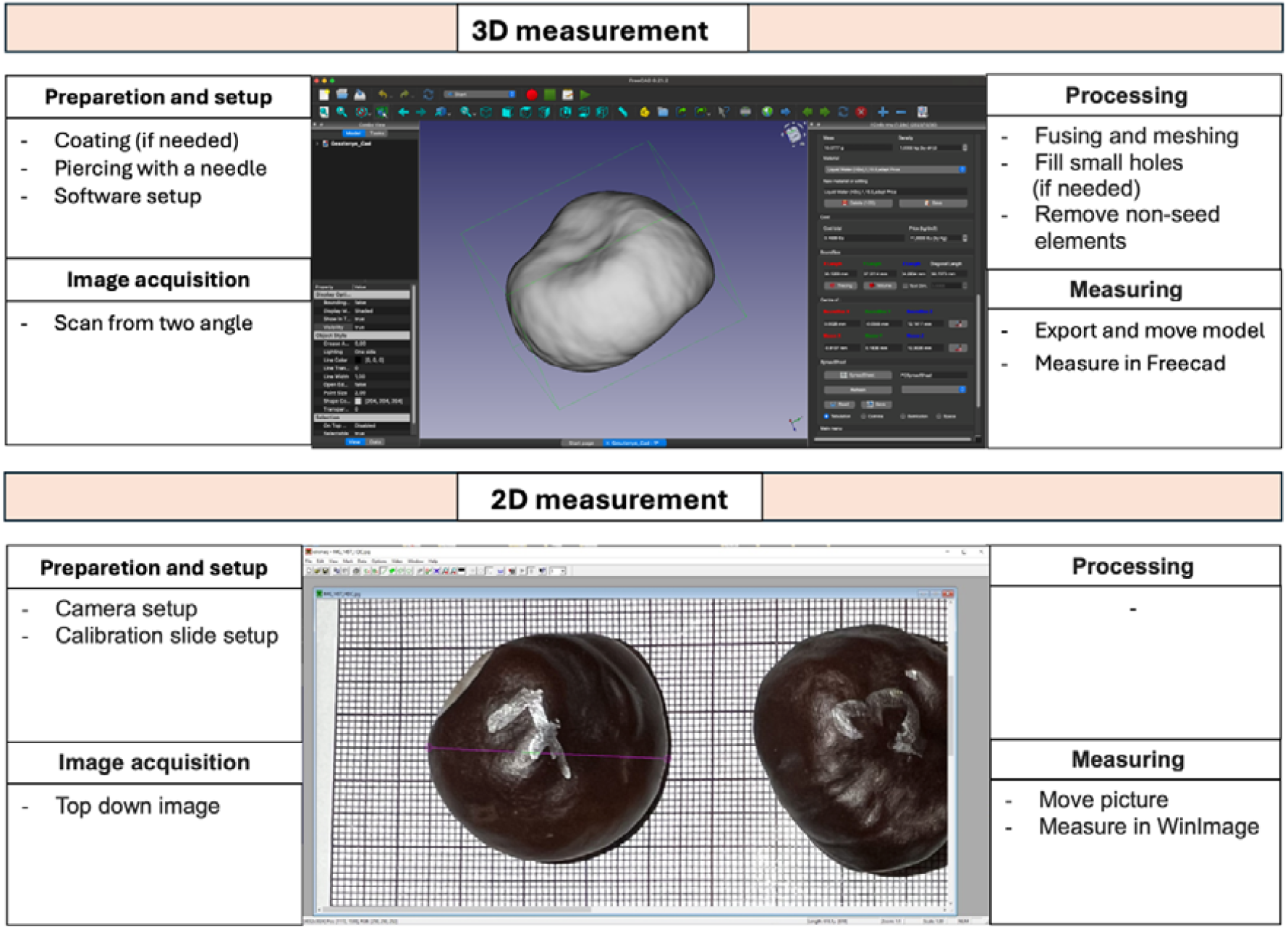
Main workflow steps for each measurement type

### 2.6 Manual measurement

Seeds length and width were also measured manually. While any digital camera will suffice, we used a Canon EOS R and a microscope calibration slide with 0.1mm precision for size reference. We analyzed images for length and width using WinImage 1.0 software (Figure 3).

### 2.7 Reproducibility of measurement

Multiple analyses were performed to determine the reproducibility: coefficient of variation (CV), paired T-test and Friedmann-test.

The overall median CV was 2.45% for 3D measurements and 3.15% for 2D measurements (Figure 4). Higher values and variations were found for Aesculus hippocastanum seeds (median: 3.06%, 1.9-5.66%), and lower values and variations were found for Quercus robur seeds (median: 1.84%, 0.8-2.66%).

**Figure 4.**
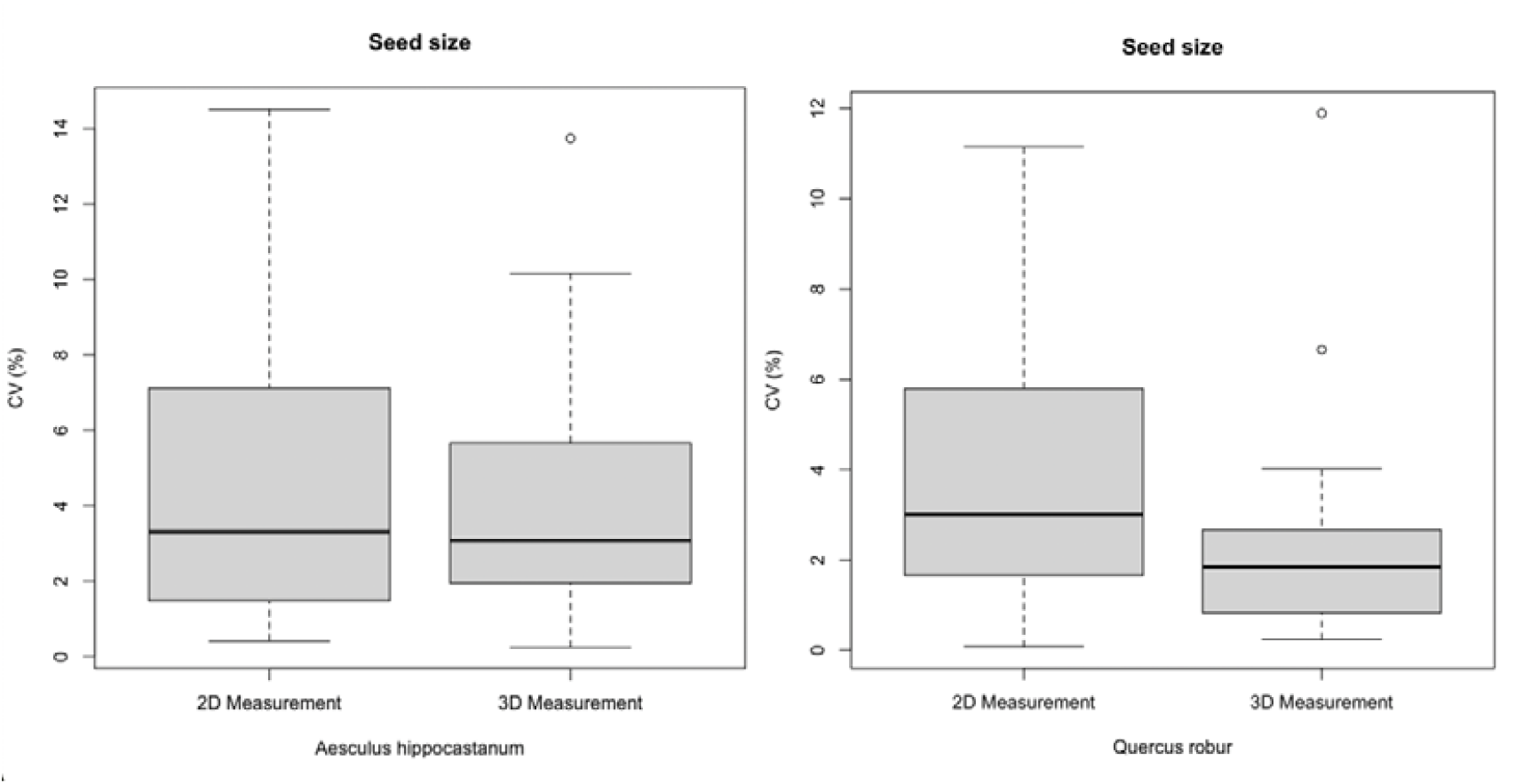
Coefficient of variation (CV) in 2D and 3D measurements

In 2D measuring, higher values and variations were found for *Aesculus hippocastanum* seeds (median: 3.30%, 1.48-7.11%) and lower were found for *Quercus robur* seeds (median: 2.99%, 1.6-5.79%).

Paired T-test showed significant differences for *Aesculus hippocastanum* seeds for both measurement group, but p-value was slightly higher for 3D (2D: t = 7.3547, df = 29, p= 4.209e-08, 3D: t = 4.6666, df = 29, p = 6.402e-05). For *Quercus robur* no significant differences were found, with higher p-value for the 2D group (2D: t = -0.30416, df = 29, p = 0.7632, 3D: t = -4.1784, df = 29, p-value = 0.000246).

Friedmann-test confirms that there is a significant difference in the overall measurements for both seed (*Aesculus hippocastanum*: χ^2^ = 48.291, df = 3, p = 1.847e-10, *Quercus robur*: χ^2^ = 31.56, df = 3, p = 6.479e-07).

The 2D (manual) measuring was done by two research team members, analyzing the same pictures taken of the seeds with the same WinImage 1.0 software. Even if both researchers try to achieve the greatest length and width of a seed, there can be a strong person-effect for 2D measurements (Ströbel et al., 2018).

It is worth emphasizing that the 3D scans were measured with a standard tool, leaving variation only possible between scans. Due these differences, we put the *Aesculus hippocastanum* seeds in a graduated cylinder to measure their volume. The cylinder diameter was 4.2 cm, with 2 ml divisions. The scanned seeds volume were calculated with Freecad (v0.21.1) software via the FCInfo (v1.28c) macro addon. The results were compared with paired T-test. It showed significant matches with the second scanning (t = 2.1626, df = 14, p = 0.04837). There were two higher differences around 1000 mm^3^ ≈ 1 ml, all others were less then 500 mm^3^ ≈ 0,5 ml. If we exclude these two differences go down greatly (t = 1.412, df = 12, p = 0.1833). This indicating that a bit training and care are necessary to create accurate 3D models, but it will remove all the observer’s errors during measuring.

### 2.8 Timing

Manual measuring could be significantly faster for trained observers than 3D scanning (Table 1). Additionally, the model processing time associated with 3D technologies is heavily dependent on the processing power of the computer and the speed of the associated storage. Those 3D files that include every step of object processing, not just the output, easily result in 100-200MB files in each case. Processing and loading times would increase significantly on older hardware. Note, that no extra time added for preparation like coating and piercing in case of 3D scanning, because during the model processing there is more than enough time to prepare the next sample.

**Table 1.** Average time required for a trained observer to complete each stage of 3D and 2D measuring.

| Stage | Duration (min) |  |
| --- | --- | --- |
|  | 3D measurement | 2D measurement |
| Image acquisition | 2:00 – 5:00 | 0:15 – 0:20 |
| Model processing | 4:00 – 5:00 | – |
| Export | 0:15 – 0:20 | 0:15 – 0:20 |

| Measure | 0:15 – 0:20 | 0:30 – 1:00 |
| --- | --- | --- |
| **Total** | 6:30 – 9:40 | 1:00 – 1:40 |

Image acquisition – Placing object in front of the 3D scanner or camera and capture it; Model processing – Fusing and Meshing the object; Export – Export and move the finished model to Freecad or move the picture from the camera to WinImage software in the computer; Measure – Measure the model or picture.

## 3. DISCUSSION

3D measurement techniques are not new to specific research areas of ecology. Several studies used 3D scanners to measure coral microfragment area or growth, where traditional methods are often destructive (Reichert et al., 2016; Quigley, 2020; Koch et al., 2021; Reichert et al., 2017). It creates a highly detailed model for small objects where precision and accuracy are key. However, it can also be used to create accurate models from animal skeletons for field identification or education (Laura et al., 2009). Our goal with this study was to promote usage for other areas where these tools are not utilized yet.

Compared to the manual methods of estimating seed size or surface area, 3D scanning has several advantages. Based on our tests, 3D measuring revealed significant differences compared to 2D measuring in length and width results, which could lead to the introduction of noise and potential bias when compiling datasets. With a bit training, 3D result for volume indicated similar result with what measured with a cylinder. Even if we consider the higher analysis time with modern hardware, the advantages are numerous. Universal file formats make result models easily transferable to any open-source or propriety software for further analysis. The high-resolution data can be used to share between researchers, to create an accurate, cost-effective and reproducible 3D prints of the seeds (Behm et al., 2018; Schtickzelle et al., 2020) It also has educational purposes as the 3D models also can help to attract more youth into to world of ecology (Das et al., 2019). Although the price for a professional level 3D scanner or printer is still very high, consumer level equipment delivers decent results in close-range applications. The above-mentioned scanner has some limitation regarding how small seed it can scan, for test we also tried to measure a couple of Vitis vinifera (*Vitaceae*). After the same preparation work and we could see the outlines in the software, but it would not allow a normal scan with Feature Tracking, because it didn’t have enough point clouds. We were not able to determine how to turn of this software limitations. Surface area calculation is a challenging task with manual measurement due to the curvation of the seeds, but 3D scanning has the capacity to provide more accurate measurements, as the above-mentioned tools would automatically calculate it from the model. It could also open new ways for seed shape analysis (Dayrell et al., 2023; Cervantes et al., 2016).

## 4. CONCLUSION

The presents study demonstrates that 3D scanning could provide a more precise, reproducible and consistent method for seed measurement, comparable to 2D methods. We described a general workflow for 3D scanning seeds with high efficiency and showed advantages and disadvantages. While 3D scanning may not be as time effective, it provides additional research opportunities with the 3D models. As with any new technology, researchers must spend some time to get familiar with it, but we believe it is likely to become a valuable tool in more and more ecologist’s toolbox.

## Acknowledgments

This research was partly supported by ÚNKP-23-4-I-DE-303

## Authors’ Contributions

Conceptualization, H.M-R., B.T.; Validation, H.M-R.; Resources, H.M-R. and T.M.; Writing – original draft, H.M-R., T.M; Writing – review & editing, B.T., D.T. and T.M.; Visualization, H.M-R. All authors have read and agreed to the published version of the manuscript.

## Notes

### Competing Interest Statement

The authors have declared no competing interest.

